# Effects of shock on perceptual learning in a virtual reality environment

**DOI:** 10.1101/2025.01.19.633815

**Authors:** John Cass, Wing Hong Fu, Yanping Li, Larissa Cahill, Gabrielle Weidemann

## Abstract

This study investigates two aspects of visual processing and perceptual learning: the impact of training on the human visual system’s ability to integrate information across the visual field and the influence of aversive electrodermal stimulation on perceptual performance in an orientation averaging task. Through a ten-day training regimen that manipulated the set-size of Gabor element arrays, we observe consistent degradation in orientation averaging performance with increasing set-size, with training leading to overall improvements in accuracy and response times, but only a marginal interaction with set-size (for response times). This suggests that training-related enhancements likely operate at a post-integration and/or decisional stage of processing rather than at an early encoding stage. The second inquiry explores the effects of aversive stimuli on perceptual learning-based orientation averaging performance, with participants exposed to ‘no shock’, ‘performance-contingent shock’ or ‘random shock’ conditions. Our results show that while performance improved across training there was no discernible effect of the shock condition on task accuracy or response times and no evidence of an interaction with set size. State anxiety levels, measured by the State-Trait Anxiety Inventory, indicated that whilst anxiety was elevated in both shock conditions, this was not associated with variations on orientation averaging performance. We also find that visual performance feedback, represented by a health bar, significantly influenced accuracy, but not response times, regardless of the presence or absence of shock. This unexpected impact of visual feedback suggests potential roles for attention and motivation in perceptual performance.

**Public significance statement:** This study enhances our understanding of how training and stress influence visual processing and perceptual learning, revealing critical insights into the mechanisms that govern how we integrate visual information. By exploring the effects of acute stress and visual feedback, our findings suggest potential applications for improving learning outcomes in various contexts, from virtual to real-world environments.

## Introduction

Modern visual display devices, including screens and head-mounted systems, can deliver an extensive amount of information to the human visual system with high spatial and temporal resolution [1]. However, not all information presented to the retina is perceptually accessible. Far from it. The ability of the human brain to concurrently extract visual information from its surrounding environment is contingent upon a multitude of sensory, attentional and contextual factors that limit perceptual resolution and/or its capacity [2-5].

One such factor is experience. In the very short-term, exposing the retina to light; whether direct or reflected from visual objects and events; can affect our ability to subsequently detect and correctly classify subsequently presented visual stimuli [6-8], a phenomenon known as *adaptation*. In general adaptation effects are transitory, with visual performance returning to baseline (pre-adaptation) levels after several seconds to several minutes, depending upon the adapting stimulus and task [9, 10] (see [11] & [12] for some longer-term examples). Adaptation effects are also passive, insofar as they generally operate independently of the observer’s active engagement or decision-making. Experience can also exert long-lasting effects. Active training on visual tasks can lead to long-lasting improvements in visual performance; a phenomenon known as *perceptual learning*. Perceptual learning enhances the efficiency with which the visual system can extract and interpret information [13, 14]. It encompasses various aspects of sensory processing, including the detection and discrimination of objects [15, 16]. In addition to enhancing sensory processing and efficiency, perceptual training can also improve efficiency in tasks requiring the deployment of attention across the visual field [17].

One task relevant to the question of how much information the visual system can perceptually extract and integrate, that is known to benefit from training, is ensemble processing. Ensemble processing refers to the ability of the visual system to extract statistical properties, such as the average or variance, from a set of individual spatially disparate visual elements composed of locally diverse images properties [18, 19]. [19] demonstrated that when subjects were instructed to report the average global orientation of a set of simultaneously presented Gabor patches with locally diverse orientation properties, global orientation thresholds improved significantly over subsequent days of testing. This perceptual learning effect they attribute to improvements in *sampling efficiency*. In this case, sampling efficiency refers to how many pieces of the available local orientation information the visual system uses to make decisions about the global (average) orientation properties of the stimulus. According to Moerel and colleagues’ interpretation [19]training increases the amount of local orientation information (Gabor patches) that participants use to make their perceptual decisions.

One study that specifically investigated the role of sampling efficiency in orientation averaging judgements found a causal role for attention [20]. Using an equivalent noise paradigm Dakin and colleagues showed that when attention is simultaneously deployed to the fovea (using a ‘tumbling T’ identification task) and to peripheral visual locations (orientation averaging task), global orientation averaging judgements become corrupted - relative to when the foveal dual task is not present - in a manner consistent with degraded sampling efficiency. [20] proposed that there are (at least) two possible computational explanations for such degradations in sampling efficiency due to attentional load: *early selection* and *late selection*. According to the early-selection account, attentionally mediated variations in sampling efficiency occur before the neural integration of local orientation information. When attentional resources are depleted or not deployed efficiently (as occurs during their dual-task), this is assumed to reduce or impede the number of local samples transmitted to higher-level integrative and/or decisional stages of processing, leading to poorer orientation averaging performance. Consequently, if attentional resources are deployed more efficiently (because of training, for example) this may have the effect of increasing the number of local samples available at higher-level integrative stages, thereby improving orientation averaging performance. By contrast, according to the *late-selection* account, attentionally mediated variations in sampling efficiency reflect variations in signal-to-noise at a level of processing following (or commensurate with) the integration of local orientation information, possibly at a decisional stage. According to this account, depleted or inefficient deployment of attentional resources to the orientation task-relevant information serves to reduce signal-to-noise, whereas efficient deployment of attention to this information improves signal-to-noise.

To determine whether improvements in orientation-averaging performance from training [19] are a consequence of improved sampling efficiency at either an *early stage* or at a *late stage* of processing, in this study we manipulate the amount of task-relevant information available to participants across experimental trials. To accomplish this, we simply vary the number of local task-relevant oriented elements in the stimulus display, i.e. *set-size*. Because the oriented elements used in our averaging task are drawn from a broad distribution of orientations centred on the mean signal, ideal performance requires perceptual analysis of all local orientation information presented within a given display. If, for example, participants were to base their decision on a subset of the available elements this would produce sub-optimal performance across trials.

Because increasing the number of elements necessarily increases the amount of information (number of samples) that participants must extract and integrate, it is predicted that orientation averaging performance will degrade with increasing set-size. If training serves to increase sampling efficiency at an early encoding stage this will increase the number of local orientation signals available at subsequent integration/decision stages. Consequently, not only does the early selection account predict overall improvements in orientation averaging performance because of training [19], but critically the slope of performance degradation, we expect to occur with increasing set-size, is also predicted to be reduced because of training. Alternatively, if training increases sampling efficiency by increasing signal-to-noise at a late, post-integration stage of processing, it is assumed that this will have no effect on the number of local samples feeding this stage of processing. Therefore, whilst overall training-related improvements in orientation-averaging performance are predicted, late-selection predicts that the *slope* with which performance is expected to degrade with set-size is not anticipated to vary because of training.

### Aversive stimuli and perceptual learning

Perceptual learning can also be affected by associative context. Numerous studies have demonstrated that pairing an otherwise neutral (conditioned) perceptual stimulus with an aversive (unconditioned) stimulus can affect subsequent perceptual performance if the pairing is contingent on the relevant perceptual dimension. This is a form of aversive Pavlovian discrimination learning where learning to anticipate a threat improves sensory discrimination performance [21]. Pairing perceptual stimuli with different outcomes has been hypothesized to increase the distinctiveness of the stimuli in a process referred to as “acquired distinctiveness” [22]. The literature on the effects of aversive Pavlovian discrimination learning on perceptual performance is mixed, with some studies showing improvements in perceptual discrimination and others, impairments. For example, Li, Howard [23] found that mirror-symmetrical pairs of odorant molecules that were otherwise perceptually indistinguishable could be rendered discriminable as a consequence of pairing one enantiomeric version of the odorant with an electrodermal shock stimulus. Similarly, Rhodes, Ruiz [24] reported that pairing visual luminance gratings with aversive loud auditory noise bursts elicited subsequent improvements in grating orientation discrimination thresholds for orientations. Conversely, Shalev, Paz [25] found that pairing aversive images, sounds and odors [26] produced subsequent feature-specific elevations (impairments) in auditory frequency and visual contrast and orientation discrimination thresholds.

A factor that may contribute to the complex pattern of results described above is stress. Aversive stimuli and environments produce a cascade of autonomic responses linked to elevations in blood cortisol, via the hypothalamic-pituitary-adrenal axis [27]. In addition to physiological changes (increased heart-rate, blood pressure, sweating, pupil dilation) and emotional responses (increased anxiety and arousal), elevated blood cortisol is linked to changes in frontally mediated cognitive and attentional functioning [28]. Elevated blood cortisol has also been found to inhibit perceptual learning [29].

To date, no study has investigated the effect of aversive conditioning and/or stress on orientation-averaging performance. There are reasons to believe that acute stressors may have a negative impact. Acute stress has been found to deplete attentional resources, impairing performance on some (but not all) visual tasks requiring attentional deployment [30-32]. It is conceivable, therefore, that stress may not only impair orientation averaging performance overall, but it may do so by degrading the visual system’s capacity to sample from multiple retinal locations. In light of the finding that an attentionally demanding dual-task impairs orientation averaging performance by reducing effective sampling efficiency [20], it is conceivable that presenting acute stressors may have similar effects.

Perceptual performance can also be modified by introducing motivational consequences for performance [33-36]. Similar to the effects of positive reinforcement, introducing aversive behavioural *consequences* for incorrect perceptual decision-making, positive punishment, a type of operant (or instrumental) conditioning, has been shown to improve performance. For example, Erickson [34] found that contingent negative outcomes (loud noises) in response to incorrect responses produced more accurate responding than noncontingent negative outcomes in a difficult positional discrimination task. Additionally, visual discrimination performance has been shown to improve as a function of punishment magnitude suggesting that the effect of punishment on performance is a consequences of motivation [33]. To investigate whether orientation averaging performance is similarly susceptible to aversive consequences we will evaluate the effects of two types of shock condition. In one condition shock will be contingent upon performance with shock probabilistically administered following incorrect orientation averaging responses (*performance*-*contingent shock condition*). In the other condition shock will be administered at random intervals throughout the testing procedure (*random shock condition*), with a similar frequency to that of the *performance*-*contingent shock condition*.

We hypothesize that the introduction of an aversive stimulus will modulate orientation-averaging performance. It is predicted that the random shock condition will impair orientation-averaging performance relative to the no shock condition as a consequence of the acute stress induced by the unpredictable presentation of the electrodermal stimulation (shock). Predictions include decreased accuracy and increased response times under conditions involving acute stress, reflecting the potential depletion of attentional resources. It is predicted that performance in the performance-contingent shock condition will be improved compared to the random shock condition because of increased motivation to perform well to avoid the aversive outcome. What is unclear is whether sampling efficiency will be affected by the presence of the aversive shock. On the one hand, if acute stress influences the early selection we anticipate degraded performance with increasing set-size in both the random shock and the performance-contingent shock conditions compared to the no shock condition. If, on the other hand, acute stress influences late selection then we would expect impaired performance in the random shock condition relative to the performance-contingent and the no shock conditions independent of set-size.

## Method

### Participants

A total of 22 participants were initially recruited for this study. Two participants withdrew from the study, one due to COVID and the other voluntarily. The remaining 20 participants (8 female, 12 male) were healthy adults (Mean age = 26.2, *SD* = 12.4 years) with normal or corrected-to-normal vision. Three participants were excluded. One of these participants did not complete the task properly, and two incorrectly received a disproportionately large number of shocks on one or more days of testing. Participants were undergraduate and graduate students recruited using convenience sampling. All participants provided informed written consent. Experimental protocols were approved by the Western Sydney University Human Research Ethics Committee and conducted in accordance with the Declaration of Helsinki. All participants were naïve to the aims of the study, and each received financial reimbursement totalling $250 AUD.

### Design

A 3 x 10 × 5 mixed design was employed. The between-subjects variable involved the type of electrodermal shock stimulus administered to the participants and consisted of three levels: (i) *no shock*; (ii) *performance-contingent shock* (electrodermal stimulation occurs after every ten incorrect answers, to reduce habituation to the electrodermal stimulation and ensure that it is experienced as being aversive [37]), and (iii) *random shock* (electrodermal stimulation administered randomly across trials based with a 3% chance of occurring on a given trial). On average, participants in the *performance-contingent* group received 10.9 shocks per day (SD = 2.3) and those in the *random shock* group received 13.1 shocks per day (SD = 3.0). *Participants* were randomly allocated to each group prior to testing. Regarding the within-subjects variables, the first was the number of training sessions. Each participant was subjected to 10 consecutive days of testing (excluding weekends). The second within-subjects variable, set-size, refers to the number of Gabor elements presented to participants on any given experimental trial (1, 2, 3, 4 or 8 Gabors). Set-size was randomly allocated across trials.

Participant accuracy and correct reaction times were measured on each experimental trial. The task was a custom-built first-person shooter task in a virtual reality environment. On each experimental trial, participants were required to undertake two distinct tasks: a primary perceptual task, in which they had to decide whether the average orientation of a Gabor array, visually overlaid on the virtual reality environment, was tilted clockwise or counter-clockwise of vertical; and a secondary task, in which the participant must register their primary task response by ‘shooting’ one of two virtual agents in the virtual reality environment. Details of both components are presented provided below.

### Materials

#### Software

The experiment involved two major software elements. One element involved the generation and presentation of the immersive virtual reality environment (D-world). It recorded all events within the D-world, including participant behaviour. The D-world is licenced software was built under licence by MultiSim© using the Unity Engine. For this experiment the D-world simulated a grassy plain in rural Switzerland (real-world location: 46.727° latitude and 12.219° longitude). All experimental parameters, virtual events, and performance data were stored as H5 files using an inbuilt logger with an upper maximum recording rate of 500 Hz. For storage purposes, new rows of data (time stamps) were written to the H5 files only when changes occurred within a given parameter. The other major software element, written in Python Script (v3.8.6), controlled experimental parameters, key events, and the capture and encoding of behavioural data.

#### Subjective Measure

To capture participants’ subjective state and trait anxiety levels we deployed the State-Trait Anxiety Inventory for Adults (STAI-AD) [38]. The STAI-AD includes 40 items (20 state-related, 20 trait-related questions), rated on a 4-point Likert scale with a range of “almost never” to “almost always”. Examples of trait anxiety items include, ‘I am content’, ‘I am a steady person’, and ‘I worry too much over something that really doesn’t matter’. State anxiety questions include, ‘I feel calm’, ‘I feel secure’, ‘I am tense’, and ‘I am worried’. STAI-AD has shown test-retest reliability coefficients that range from .65 to .75 over a two-month interval and an internal consistency coefficient for the scale of .86 to .95 [38]. The STAI-AD surveys were presented to participants via a Qualtrics weblink on the experimenter personal tablet or laptop device and participants’ interpupillary distance was measured using the ‘GlassesOn’ application [39] via the experimenter’s personal smartphone.

#### Hardware

The experiment was run on two separate computers using Windows 10 operating systems. The first computer, responsible for the D-World and recording behavioural data (H5 files), used an Intel(R) Core(TM) i7-9700K processor running NVIDIA GeForce GTX 2060 graphics card. The second computer, linked via ethernet to the first computer, was dedicated to presenting the stimuli (D-World) to the Oculus Rift S and had an Intel(R) Core(TM) i7-9700K processor with NVIDIA GeForce GTX 2070 SUPER graphics card. Audio was presented via headphones (wired Philips/Gibson Innovation NL 5616LZ-400 SFH4).

##### Virtual Reality Headset

Participants viewed the stimuli binocularly via a consumer release Oculus Rift S with two fast-switch LCD panels with a total resolution set at 2560 × 1440 (1280 × 1440 pixels per eye) and a refresh rate of 80 Hz. The interpupillary distance of the Oculus Rift S has a fixed physical distance of 64 mm. However, the Oculus Rift S features interpupillary distance adjustment via software. The interpupillary distance of our participants ranged between 58 mm and 69 mm. The viewing distance between the participants’ eyes and the Oculus’ lens varied between participants, but all were within the range of its default adjustment distances. Participants had approximate horizontal and vertical fields of view of 88° and 94° respectively.

##### Virtual Reality Controller

A custom-built rifle ‘prop’ with corresponding spatial dimensions to the virtual rifle was created for this experiment (see **Figure 1a**). This rifle prop consisted of a 610 mm long metal rail with a 3D printed gun stock (length = 175 mm) wrapped with foam and duct tape for comfort attached to one end. A small angled ‘V’-shaped cut was made further along (90 mm) the metal rail that housed a standard left-handed Oculus Touch (second generation) controller utilised by participants to shoot the virtual rifle which records movement with six degrees of freedom. A small metal picatinny rail was positioned 260 mm along the main metal rail to allow a 3D printed vertical foregrip (radius ∼100 mm) to be attached. Finally, an orange 90 mm (diameter) stress ball was mounted at the end of the metal rail for safety. The custom-built rifle prop weighed 988 grams.

**Figure 1.**
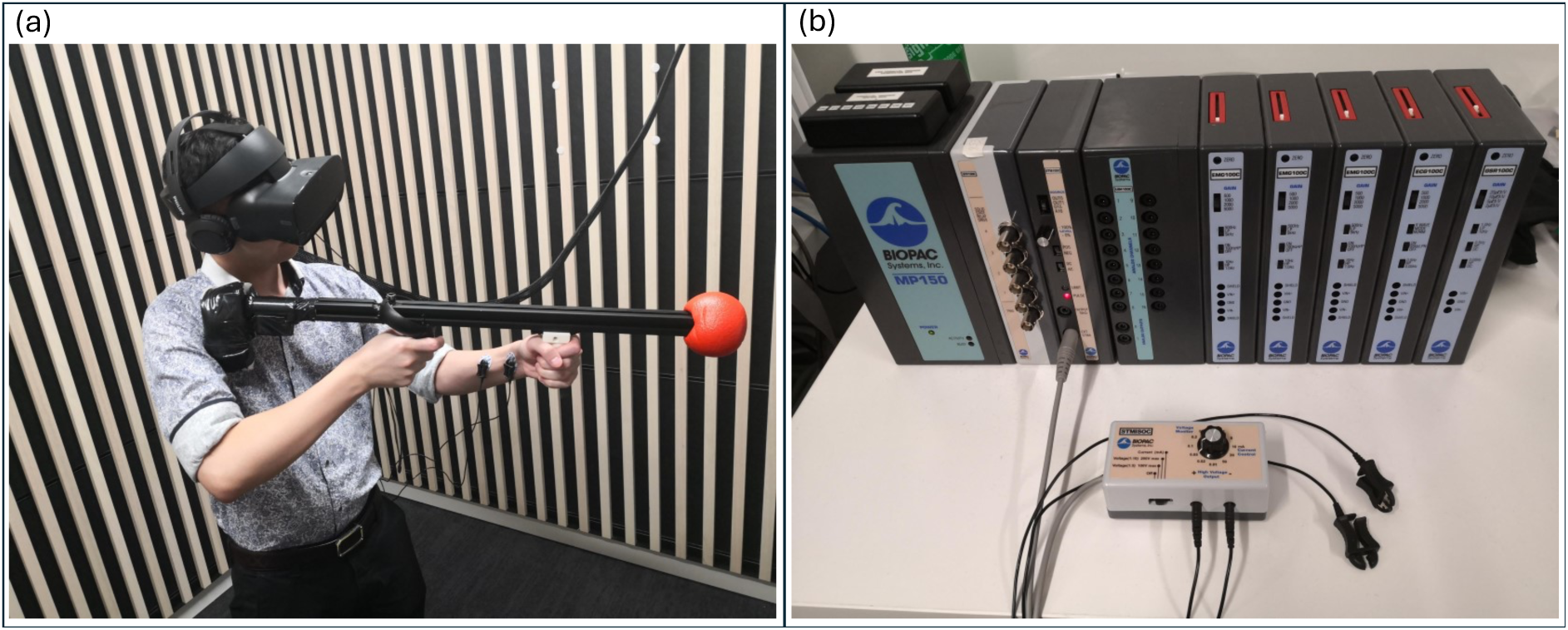
Experimental apparatus and electrodermal stimulus set-up. *Note:* Panel (a) shows a participant wearing the VR Oculus headset and headphones holding the controller-mounted custom rifle prop with electrodes applied to the non-dominant forearm. Panel (b) shows the Biopac MP-150 signal conditioning module and STMISOC electrodermal stimulation device.

##### Electrodermal Stimulation

A Biopac MP-150 was used to trigger a 100-ms electrodermal stimulation implemented with a Linear Isolated Stimulator (Stmisoc, BIOPAC Systems, Inc., Goleta, USA) to participants in the shock conditions (see **Figure 1b**). An amplifier set to “D15 (pulse)” was used as the input source, generating single positive bipolar pulses. In accordance with ethics, participants each self-selected the amperage settings, ranging from a minimum of 10 to a maximum of 100 microamps. Participants selected values ranging from 30-90 microamps (mean = 62.71, SD = 14.32).

### Stimuli

#### Specifications for Manoeuvring in the Immersive Virtual Reality environment

When entering the D-World virtual environment, participants occupied a stationary position inside the virtual world. Participant height and head movements, tracked via in-built sensors in the VR headset, were closely approximated in the D-world. Two degrees of freedom, typically expected in virtual reality environments; surge (forward/backward) and sway (left/right) (e.g., taking a step); were not available to participants in our experiment. Participants could rotationally manoeuvre their field of view across three degrees of freedom (pitch, yaw, and roll) to enable omnidirectional viewpoint shifts. Variations in height (up/down movements) were also available to the participant. After entering the D-World participants were handed the custom-built rifle prop linked to a virtual replication of a EF88 AUS Steyr rifle (see **Figure 1a**). The rifle movement had six degrees of freedom and was tracked via the left-handed Oculus Touch controller mounted beneath and near the centre of the custom-built rifle prop. Shooting was achieved by ‘pulling’ the Oculus Touch controller’s trigger.

**Figure 2.**
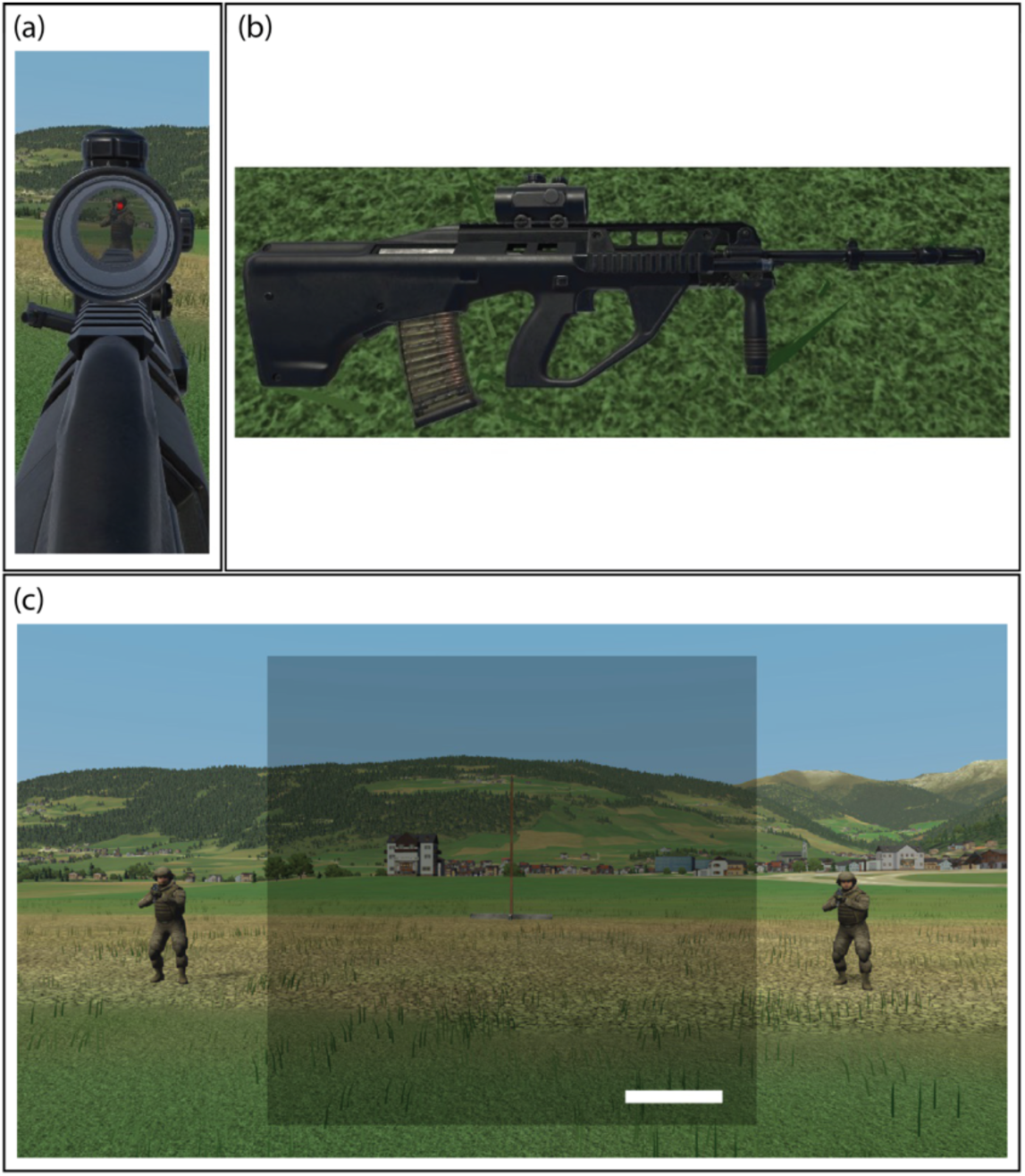
First-person perspective elements of the VR D-world environment. *Note:* Panel (a) view through the virtual rifle scope, with the red dot overlay used for aiming. Panel (b) virtual rifle laid upon the ground in the virtual environment. Panel (c) viewer perspective of the virtual environment centred upon the virtual post located between the two virtual agents. The dark semi-transparent square at the centre of the viewer’s field of view simulates a heads-up display and was fixed to the viewer’s head co-ordinates. Gabor elements appeared within this square. The white rectangle in the lower right corner was a health-bar whose length reduced by 10% following each ‘incorrect’ trial, returning to full health after reaching zero health. The health-bar provided performance feedback to all participants regardless of their assignment to different ‘shock conditions’ (see below).

#### ‘D-World’ VR Environment

Upon entering the D-World a virtual pole was positioned directly in front of the participant. On either side of the pole (a distance of 709 pixels from the centre of the pole) were two ‘soldier’ agents (98 × 256 pixels) facing the participant. Both agents were visually identical. On each experimental trial one of these agents was randomly allocated to be ‘hostile’, and the other ‘friendly’. These designations refer to the fact that on a given trial, the ‘hostile’ agent would shoot the participant if the participant failed to shoot them first. The only information available to the participant as to which agent was hostile was provided by the average orientation of the Gabor patch array presented on each trial (see below). Both the hostile and the friendly agents remained at these locations throughout the experiment and remained motionless until a ‘hostile shooting’ or ‘agent death’ animation was initiated. A transparent dark grey square (1024 × 1024 pixels) fixed to the participants’ head co-ordinates was presented throughout the experiment. This transparent dark grey square contained the orientation averaging Gabor stimuli and a small white health bar (205 × 28 pixels) located in the lower right corner.

#### Orientation Averaging Stimuli

The orientation averaging task consisted of an array of 1, 2, 3, 4 or 8 Gabor patches equidistantly positioned on an invisible annulus with a radius of 400 pixels (∼25 degrees of visual angle (d.v.a.) centred on the transparent dark grey square. The polar angle of the Gabor(s) was randomised across trials. Each Gabor patch was composed of a sinusoidal luminance grating (0.8-1.6 cycles per degree of visual angle assuming standard eye to screen viewing distance range of 2.5 – 5 cm) multiplied by a circular Gaussian envelope with a full width of 83 pixels). On each experimental trial, the global mean orientation of the Gabor array was either 5 degrees clockwise or counter-clockwise from vertical. The local orientations of each Gabor patch within arrays containing more than a single Gabor patch (set-sizes > 1) were drawn from a Gaussian distribution with a standard deviation of 20 degrees. To ensure that the mean global orientation offset of the distribution was exactly 5 degrees clockwise or counter-clockwise from vertical, the orientation of the final Gabor patch calculated within a given array was adjusted. **Figure 3** shows an example for each of the five different set sizes conditions used in the orientation averaging stimuli.

**Figure 3.**
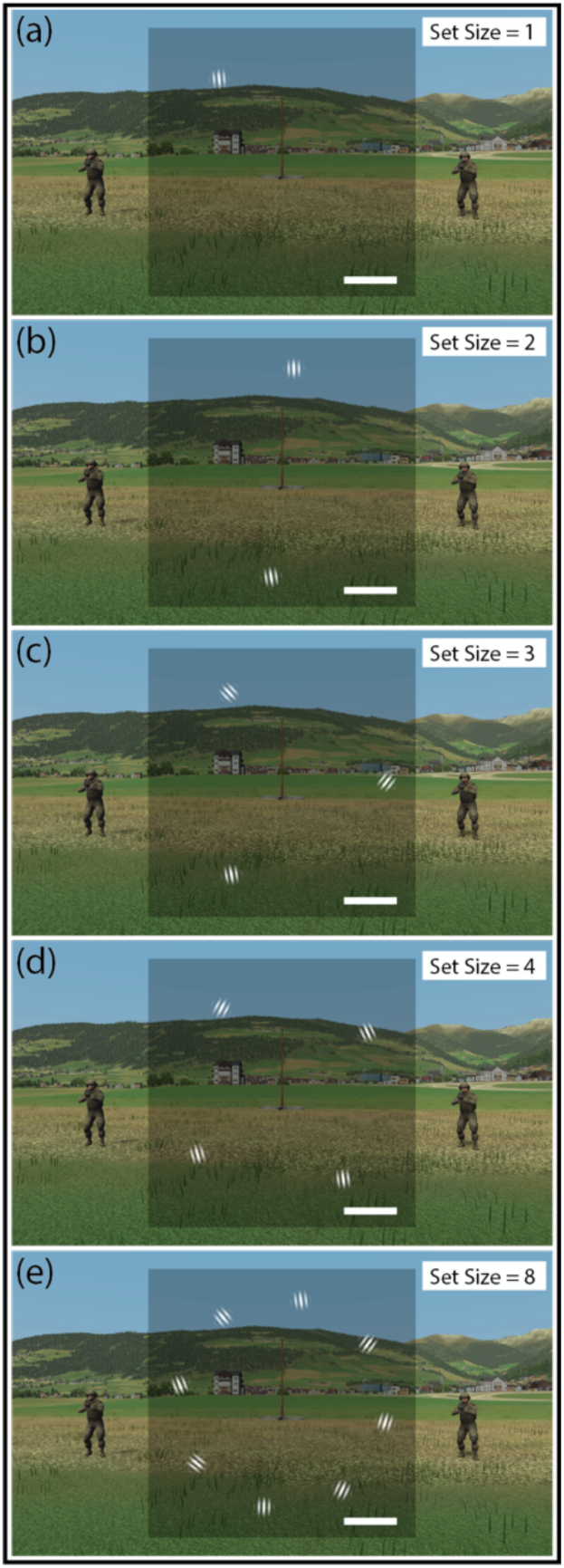
Orientation averaging stimuli. *Note.* Gabor stimuli of varying set-sizes (1, 2, 3, 4 and 8; illustrated in panels a, b, c, d, and e respectively) as viewed in the VR environment. Note on each trial the average orientation was plus or minus 5° from vertical. Images all show Gabors tilted -5° from vertical on average, signifying that the entity on the left is the enemy.

#### Time-course of orientation averaging and first-person shooter task

Each trial began with 500 ms delay where the computer systems generated the settings (i.e., Gabor distribution, enemy location, etc) and communicated with one another. Subsequently, the transparent dark grey square appeared to the participant followed by the Gabor patch array 500 ms later. The square and Gabor patches remained visible throughout the trial until the participant either shot one of the virtual soldier entities or were themselves shot by the enemy soldier. On each trial, the appearance of the Gabor patch array defined the starting point for reaction time calculations. Participants were provided with an exposure duration of 2500 ms to extract the average orientation from the Gabor patch array. Participants were instructed to aim and discharge their virtual rifle in the direction of the soldier signified by the average Gabor orientation (-5° enemy on the left, +5° enemy on the right). The positions of the two soldiers were the same on every trial, while which was ‘friend’, and which was ‘enemy’ varied randomly from trial to trial. The size of the target area to register shooting of a soldier was generous and extended beyond the soldier to ensure that left or right direction of shooting but not the accuracy of aim was the important determinant of task performance. Discharging the rifle and shooting a given soldier was relatively easy task, so shooting precision was not a factor in the experiment. After shooting one of the soldiers, the system would generate an ‘agent death’ animation depicting the soldier falling to the ground. In cases where the participant failed to shoot the enemy soldier within the allotted 2500 ms following the appearance of the Gabor array, the system generated an animation showing the ‘hostile shooting’ of the participant.

Following ‘incorrect’ trials (enemy shoots participant or participant shoots the friendly soldier) the health bar dropped by 10%. For all participants (*no shock, random shock and performance contingent* groups) this reduction in the visual representation of health acts as visual feedback about whether they successfully shot the hostile agent. For participants assigned to the *performance-contingent* shock condition, this health reduction signified that they were now 10% closer to receiving electrodermal stimulation. Participants in the *performance-contingent shock condition* received electrodermal stimulation only when their health bar reached 0%. For participants in the *random shock* condition, the health bar status was unpredictive of shock, receiving a 3% chance of receiving electrodermal stimulation at the end of a given trial. This value was chosen to approximate the number of number shocks received by participants in the *performance-contingent shock condition*, as determined in a pilot study. At the conclusion of each trial the Gabors disappeared, indicating the beginning of a new trial.

### Procedure

An identical testing procedure was employed for each of the ten consecutive days (except weekends). The only exception to this was the first training session, which began by providing participants with information about the task and obtaining informed written consent. Participants were then randomly assigned into a condition (*no shock* n=6; *random-shock* n=6; *performance-contingent shock* n=8). We then measured the participants’ interpupillary distance using the ‘GlassesOn’ smartphone application (6over6 Vision LTD, 2022) and teaching them how to wear the Oculus Rift S. The orientation averaging and shooting tasks were explained to each participant using a booklet with images of the stimuli. Each participant engaged in a practice session which consisted of several practice rounds that continued until the experimenter was confident that the participants understood and could perform the task competently. Typically, twenty trials were required to achieve competence.

Participants completed the STAI-ID prior to each training session. We refer to this as the pre-test. Following this, participants in the two shock conditions had two electrodes attached to their non-dominant forearm (the hand holding the foregrip – see **Figure 1a**). These electrodes were connected to the electrical stimulation device (**Figure 1b**). On each day of testing each participant performed a ‘work-up’ procedure which began by exposing the participant to a minimal electrodermal stimulation (∼1000 ms) of ten volts – manually controlled by the experimenter. Subsequently, the experimenter asked the participant themselves to incrementally increase the voltage followed by manual electrical stimulation until they felt that electrodermal stimulations reached a level that is “subjectively uncomfortable but not painful”. Participants assigned to the *no shock* condition never received any electrodermal stimulation and did not have any electrodes placed on their forearm. The voltages selected by the various participants average voltages across the ten training sessions ranged from 30 to 83 volts for those in the *performance-contingent shock* group and 45 to 77 volts for the *random shock* group. Participants then refitted the Oculus Rift S and stood upright. Once participants confirmed they were comfortable and ready, they undertook the experiment. This consisted of two blocks of the orientation averaging/first-person shooter task with a short five to ten minutes break between each block. Each block consisted of 210 orientation averaging trials which were evenly split between clockwise and counter-clockwise tilts of variable set sizes (i.e. 42 trials per set size). Finally, each training session ended with participants completing the STAI-ID again as the post-test for that training session.

This study was not preregistered. All data are available upon request.

## Results

In this study we assessed two measures of behavioural performance, accuracy and correct response times. Accuracy was indexed as the proportion of trials in which the participant correctly identified and shot the enemy agent prior to being shot by the enemy agent. Trials in which the participant shot the friendly agent or were shot by the enemy agent were considered incorrect trials. Response times refer to the time between the appearance of the diagnostic Gabor elements and the participant shooting the enemy agent. Only ‘correct’ trials (those where the participant shot the hostile agent before the hostile agent was able to shoot) were used in the calculation of response times.

The overall effects of set-size and training session on accuracy and correct reaction times orientation-averaging/shooting task averaged across participants in each of the three shock conditions are shown in **Figure 4**. Visual inspection of this figure reveals several trends. Most notable are the effects of training session, with performance generally improving (accuracy increasing (top row **Figure 4a-e**), response times decreasing (bottom row **Figure 4f-h**)) with subsequent days of testing. With regard to set-size (columns **Figure 4a-e**), one can observe overall reductions in accuracy and increases in response times with increasing set-size. The effects of our various shock conditions (coloured curves) are less obvious from visual inspection.

**Figure 4.**
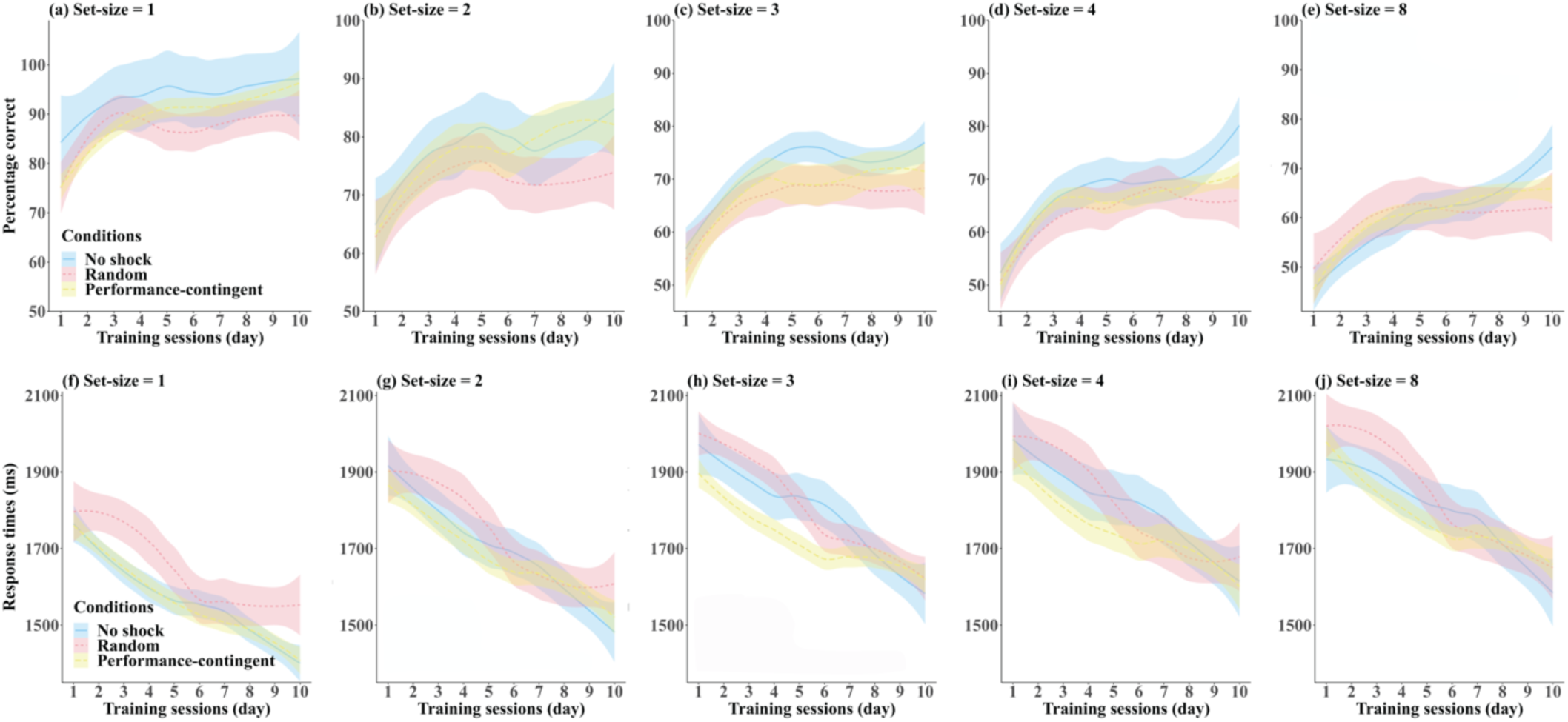
Accuracy and response times. *Note:* Effects of training session and set-size on participant-averaged accuracy (a-e) and response times (f-g) in each of the three shock condition training groups: no shock (blue), random shock (red) and performance-contingent shock (yellow). Shaded regions are between-subjects standard errors.

To statistically assess the main and interactive effects of training and shock condition we applied growth curve analysis [40]. Growth curve analysis can be used to analyse performance changes in longitudinal studies, so we employed this technique to model the effects of training in our orientation-averaging task. This was done using the *lmer* function from the package *lmerTest* [41] in *R* version 4.2.1 [42]. Given that several of the curves in **Figure 5** had at least a single inflection, growth curve data were modelled with up to second-order orthogonal polynomials, which were assessed by three terms in the fitted model: the intercept describes the mean values, the linear term captures the negative or positive slope of the curve over time, and the second-order quadratic term signifies degree of inflection in curve complexity, i.e., the depth of any peak or valley in the training curve over time. Differences in at least one of the three terms (intercept, linear, and quadratic) must be significant to indicate a reliable growth curve difference between the three shock conditions groups.

**Figure 5.**
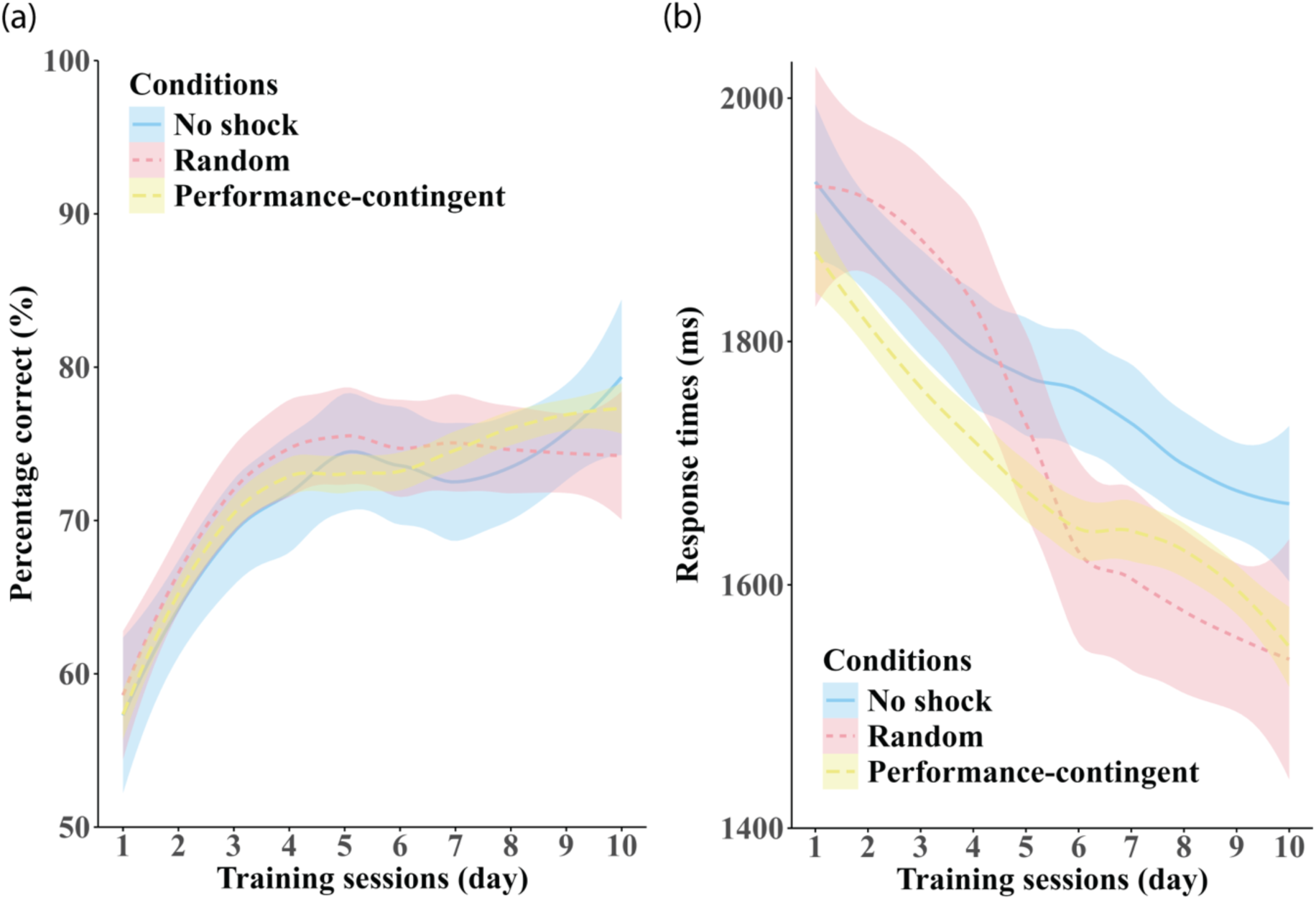
Effects of training session. *Note:* Panel (a) shows on participant-averaged accuracy and panel (b) shows response times averaged across set-size in each of the three shock-conditions: no shock (blue), random shock (red) and performance-contingent shock (yellow). Shaded regions are between-subjects standard errors.

### Training effects

Whether participants performed differently across training sessions was assessed by fitting linear mixed-effect models with shock condition and training session as fixed effects and set-size and participants as random effects. The Kenward-Roger degrees of freedom approximation [43] was used to calculate *p* values for the fixed-effect factors, and the *ANOVA* function from package *car* [44] was used to calculate *F*. Pairwise comparisons were conducted using the *lsmeans* package [45] in *R* when necessary. Linear mixed-effect ANOVAs on our performance measures were undertaken using Type-II Wald F tests with Kenward-Roger degrees of freedom.

### Accuracy

Linear-mixed effects modelling revealed no main effect of training shock condition on accuracy, *F*(2, 13) = 0.004, *p* = .996. Averaging across shock condition, we observe a significant main effect of training session, *F*(9, 754.02) = 54.639, *p* < .0001, with improvements in accuracy observed across training sessions. No interaction was observed between training session and shock condition, *F*(18, 754.02) = 1.04, *p* = .343. Planned comparisons of accuracy obtained between subsequent days of testing (collapsed across shock condition) showed improvements in performance only occurred with respect to the first day of testing (day 1 vs days, 2-10, all *p*-values < .001). No significant differences in accuracy were observed across subsequent consecutive days of testing (all *p-*values > .05).

### Response times

Like accuracy, linear mixed-effects modelling found no evidence for a main effect of shock condition on response times *F*(2, 13) = 0.453, *p* = .646. Also similar to our accuracy measure, averaging across shock condition, we observe a significant main effect of training session on response times, *F*(9, 745.01) = 99.249, *p* < .0001, with significant improvements (reductions) in response times across training sessions. Planned contrasts comparing response times obtained on subsequent days of testing (collapsed across shock condition) showed significant reductions only occurred between days 4-5, t(745) = 6.180, *p* < .001, and 6-7, t(745) = 3.566, *p* = .007. All other *p*-values > .05.

Unlike accuracy, we observe a significant interaction between training session and shock condition on response times, *F*(18, 745.01) = 5.181, *p* < .001. To further understand the nature of this interaction between the effects of training session and shock condition we first compared the relative fits of the three training-related growth curve models (intercept, linear, and quadratic) for each pair of our three training condition groups. No significant differences were observed between any of our shock condition groups in any version of our training-related response time growth curve model (all *p*-values > .05). We next examined whether mean response times improved (decreased) across days of testing both *between* and *within* each shock condition group. No differences in response times were observed *between* the different shock conditions on any given day of testing (all *p*-values >.05). For comparisons between each day and all subsequent days of testing *within* each shock condition group we observe the following distinctive patterns across the various shock conditions. Notably, whereas robust improvements in response times were observed following the first day of testing (on all subsequent days) in all shock conditions (all *p*-values < .05), for response times significant reductions in response times were not observed until day 3, 4 and 5 in the *performance-contingent*, *no-shock* and *random-shock* conditions respectively. Relative to day 3, whereas improvements were observed from day 7 in the *no-shock* and *performance-contingent shock* conditions, in the *random shock* condition faster responses were observed from day 5. Relative to day 4, whereas improvements in response times were observed from day 8 in the *no-shock* condition and days 7, 9 and 10 in the *performance-contingent shock* condition, improvements were observed from day 7 in *random shock* condition. Relative to day 5, whereas no subsequent improvements were observed on in either the *no-shock* or the *performance-contingent shock* conditions, improvements were observed on all subsequent days of testing in the *random shock* condition and on day 10 in the *performance-contingent* condition. Finally, whereas no subsequent improvements in response times were observed in the *no shock or random conditions*, significant improvements were observed in both the *performance-contingent* condition from days 8-10.

### Set-size effects

**Figure 6.**
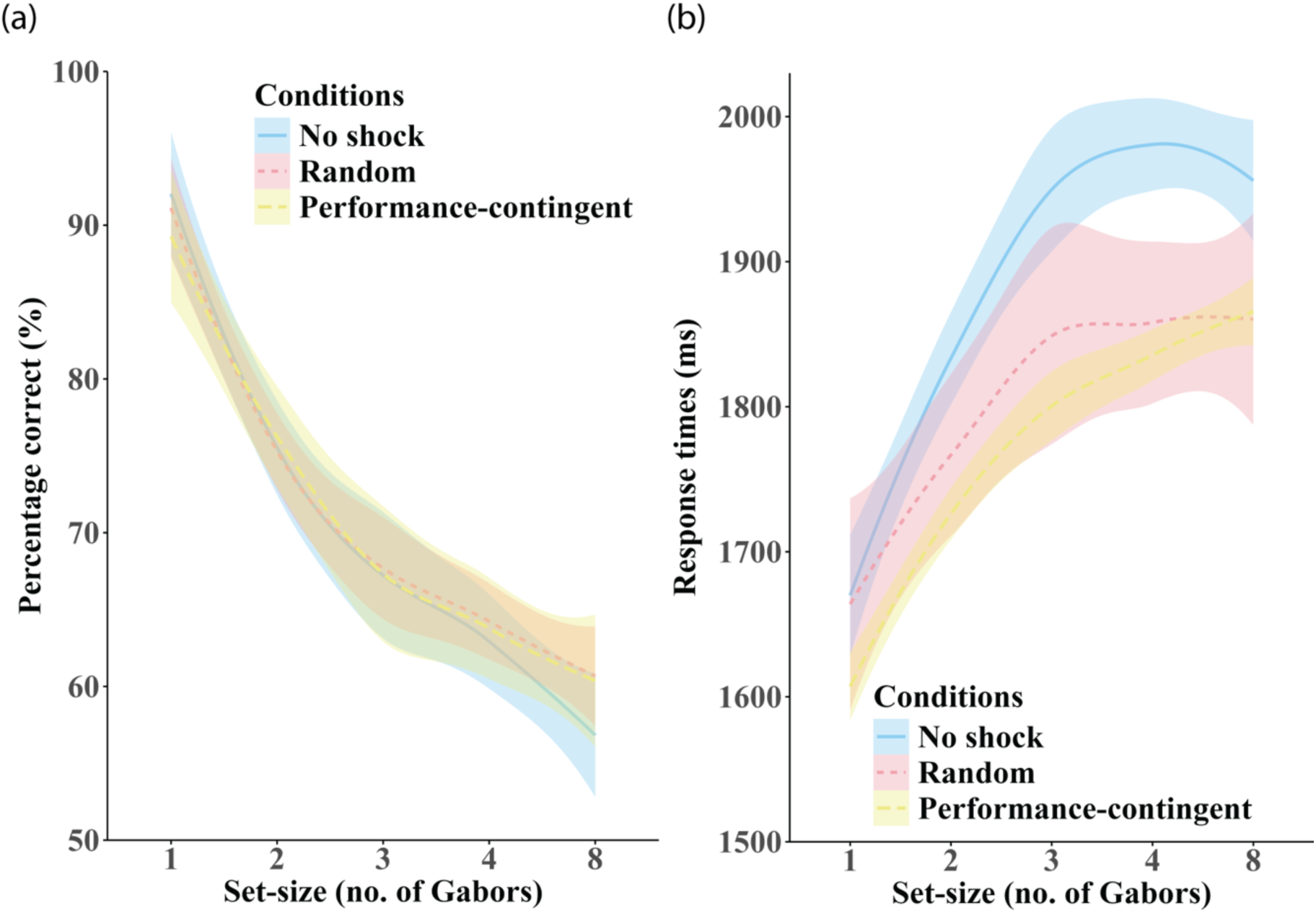
Effects of set-size. *Note*: Panel (a) shows participant-averaged accuracy and panel (b) shows response times averaged across training session in each of the three shock-conditions: no shock (blue), random shock (red) and performance-contingent shock (yellow). Shaded regions are between-subjects standard errors.

### Accuracy

To statistically assess the relationship between set size and accuracy we applied growth curve analyses using set-size and shock regime as fixed effects with training and participants as random effects. Linear-mixed effects modelling revealed a significant main effect of set-size on accuracy, *F*(4, 755) = 484.951, *p* < .0001, with decrements in accuracy linked to increasing set-size. Planned contrasts between all set-size combinations indicated that increasing set-size universally resulted in decrements in accuracy (all *p*-values < .0001, except set-size 3 vs 4, *p*-value = .006). A significant interaction was observed between set-size and shock condition, *F*(8, 755) = 2.186, *p* = .026. To understand the nature of this interaction we next compared the relative fits of the three set-size-related growth curve models (intercept, linear, and quadratic) for each pair of our three training regime groups. No significant differences were observed between any of our shock condition groups in any of the three models tested (all *p*-values > .05). To further deconstruct the interaction, we next performed a series of pairwise contrasts (with Bonferonni adjustment) to examine differences in accuracy across set-size both *within* each shock condition group and *between* shock condition group at each set-size tested.

Comparing accuracy obtained *within* each shock condition, with several exceptions (see below) that increasing set-size led to poorer accuracy (all *p*-values < .005). Exceptions to this are as follows: *No-shock* (set-size 3 vs 4, *p* = .624)); *Random shock* (set-size 3 vs 4, *p* = .868); *Performance-contingent shock* (set-size 3 vs 4, *p* = .871). No pairwise differences in accuracy were observed *between* shock conditions for any given set-size (all *p*-values > .05).

### Response times

To statistically assess the effect of set-size on response times we applied linear mixed-effect modelling with shock condition and set-size as fixed effects, and training session and participants as random effects. We observed a significant main effect of set-size on response times, *F*(4, 755) = 118.530, *p* < .0001, with longer response times associated with increasing set-size. Planned pairwise contrasts between all set-size combinations (collapsed across shock condition) indicated that increasing set-size resulted in significantly longer response times (all *p*-values < .0001), with the following exceptions: 3 vs 4, 3 vs 8 & 4 vs 8 (*p*-values > .05), indicating that response times did not increases for set-sizes greater than 3.

A significant interaction was observed between set-size and shock condition, *F*(8, 755) = 2.423, *p* = .013. To understand the nature of this interaction we statistically evaluated the relative fits of our three growth curve models. No differences in fits were observed between any training regime using and version of our growth curve model (intercept, linear, or quadratic) (all *p*-values > .05). To further investigate the nature of the observed interaction between set-size and shock condition on response times we conducted a series of contrasts, comparing response times across set-size both *within* each shock condition group and *between* shock condition measured at each set-size tested. With the following exceptions (see below) we find that increasing set-size led to longer response times (all *p*-values < .05). Exceptions are as follows: *No-shock* (set-size 3 vs 4, *p* = .999; 4 vs 8, *p* = .999); *Random shock* (set-size 3 vs 4, *p* = .996; 4 vs 8, *p* = .999); *Performance-contingent shock* set-size 2 vs 3, *p* = .130; 3 vs 4, *p* = .981; 4 vs 8, *p* = .743). No pairwise differences in response times were observed *between* shock conditions for any given set-size (all *p*-values > .05).

### Training x set-size effects

To statistically assess the existence of any the main and interactive effects training session and set-size on accuracy and response times we applied growth curve analyses separately for accuracy and response time data using set-size and training session as fixed effects participants as random effects. Consistent with the growth curve analyses described above, linear-mixed effect modelling revealed significant main effects of training session for both accuracy, *F*(9, 727.01) = 53.634, *p* < .0001, and response times, *F*(9, 727.01) = 86.607, *p* < .0001. Similarly, main effects of set-size were observed for both accuracy, *F*(4, 727.00) = 471.307, *p* < .0001, and response times, *F*(4, 727.00) = 111.968, *p* < .0001. No interaction between training session and set-size was observed for either accuracy, *F*(36, 727.00) = 0.660, *p* = .9441, or response times, *F*(36, 727.00) = 0.125, *p* = 1.0.

### Health-bar feedback

For the next series of analyses, we investigated whether performance feedback visible to participants via the ‘health bar’ (present throughout each training session) affected participant performance (see Methods for information on health bar performance feedback).

**Figure 7a & b** shows the relationship between health-bar level (low (≤ 20%), medium (21 – 80%) and high values (>80%)) on average accuracy and response times, respectively, in each of the three group training conditions. We built two sets of linear-effects models, one set for accuracy, and one for response times, with health-bar level and shock condition as fixed factors and participant as a random factor. Linear mixed effects modelling yielded a significant main effect of health-bar index on accuracy, *F*(2, 922) = 497.380, *p* < .0001. Contrasts comparing accuracy obtained at each health-bar level (low vs medium, low vs high, medium vs high) indicate that accuracy improved with increasing visual health-bar level (all *p*-values < .0001). Health-bar level had no effect on response times *F*(2, 922) = 0.310, *p* > .05. Bonferroni-adjusted pairwise comparisons showed that accuracy increased with increasing health-bar status in each of the three shock conditions (all *p*-values < .0001). No main effect of shock condition was observed for either accuracy, *F*(2, 922) = 0.001, *p* > .05, or response times, *F*(2, 922) = 0.475, *p* = .632. No significant interactions between health-bar level and shock condition were observed for either accuracy, *F*(2, 922) = 0.326, *p* > .05, or response times, *F*(2, 922) = 0.299, *p* > .05.

**Figure 7.**
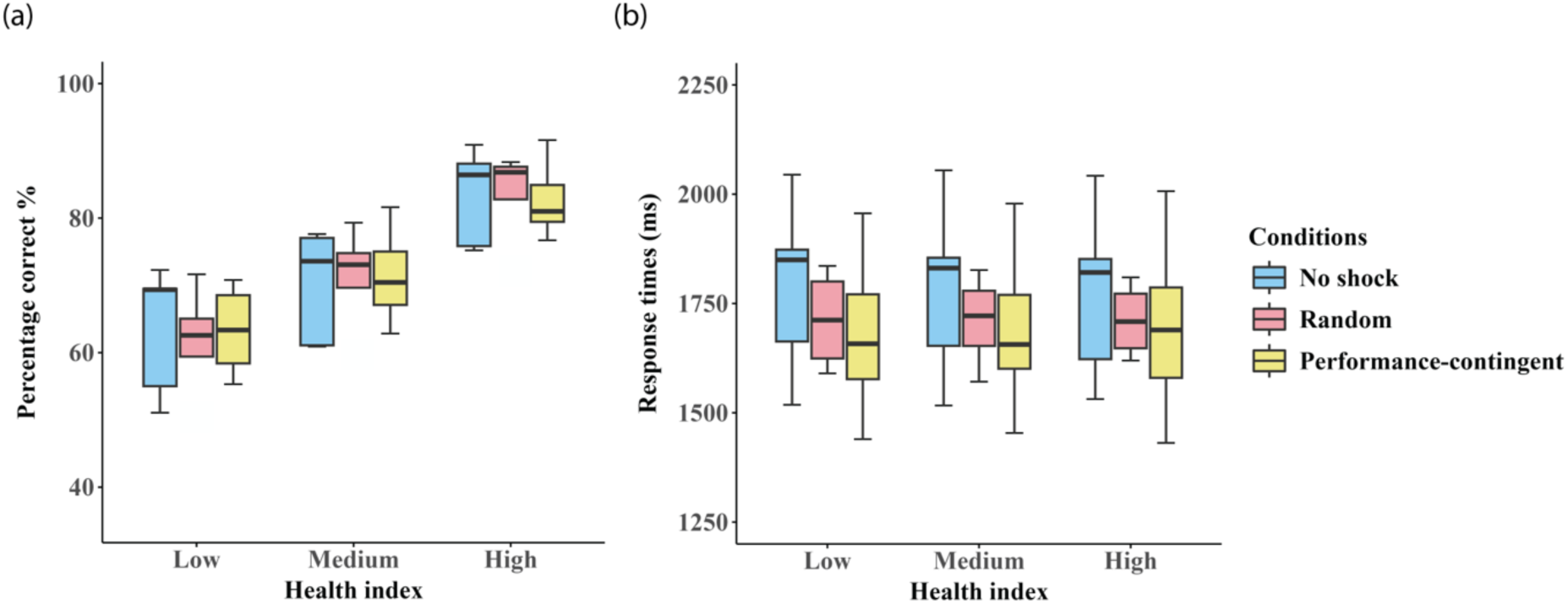
Effects of visual health-bar status. *Note:* Box plots showing the effect of visual health-bar performance feedback (low (≤ 20%), medium (21 – 80%) and high values (>80%)) on average accuracy are shown in panel (a) and response times in panel (b) in each of the three shock groups: no shock (blue), random shock (red), and performance-contingent shock (yellow). Error bars represent between subject standard errors.

The health-bar provides performance feedback to the participant, dropping 10% following each incorrect trial, and refreshing after each 10^th^ incorrect trial. It is worth noting that the health bar status is predictive and consequential only for participants in the *performance-contingent* shock group who received a shock following nine previous errors. The health bar level is completely inconsequential (i.e. unpredictive of the of a shock stimulus) for participants in both the *no-shock* and *random shock* groups. To determine whether there exist additional effects of anticipating and/or receiving a physical shock, we conducted additional analyses evaluating the effects on performance (accuracy and response times) on the five trials immediately preceding and the five trials immediately succeeding a physical shock (**Figure 8**). Given that shock was only presented to participants in the *random* and *performance-contingent shock* conditions, participants in the *no-shock* condition were omitted from these analyses.

**Figure 8.**
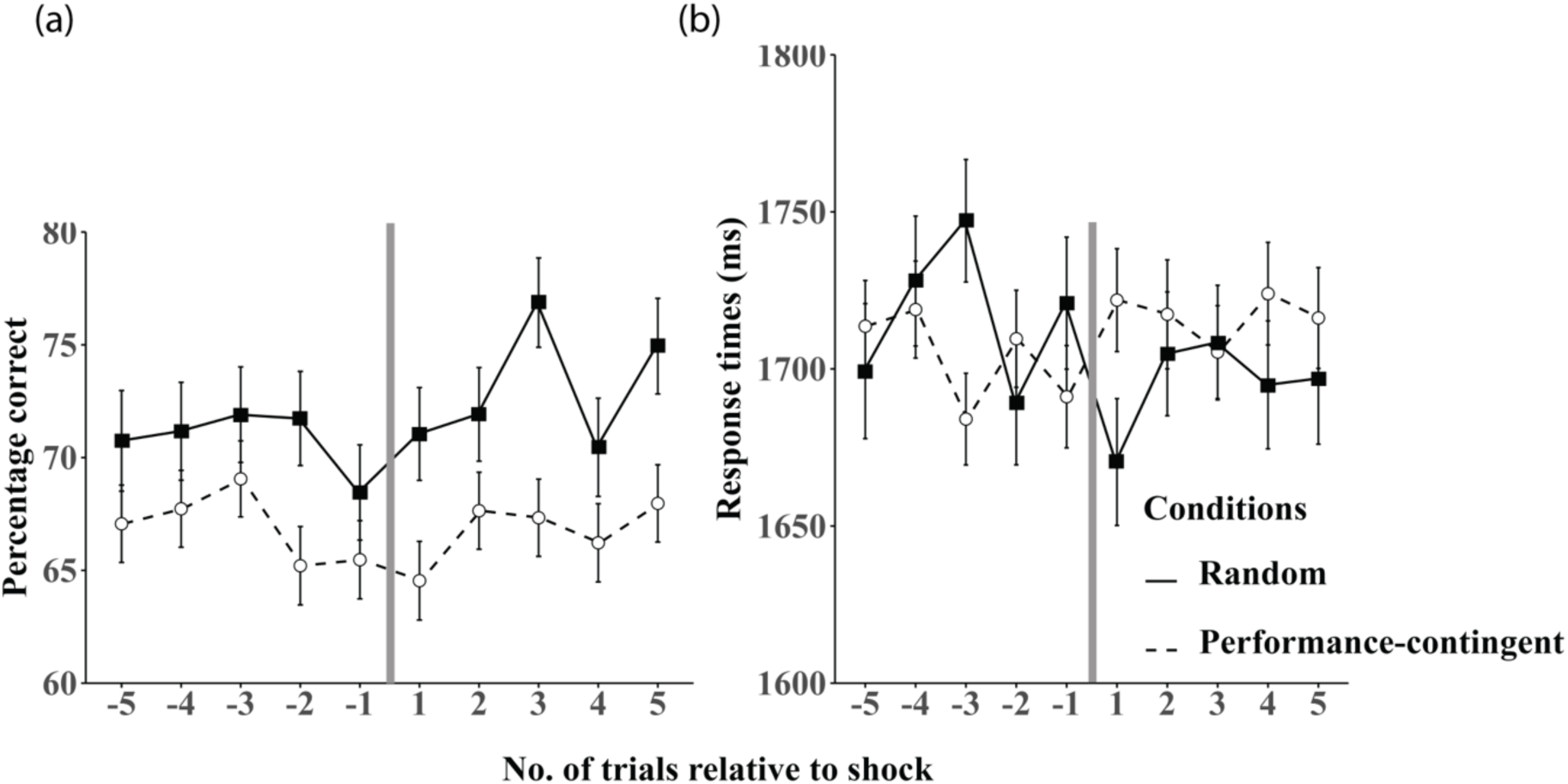
Performance relative to shock delivery. *Note:* Mean accuracy on the five trials preceding and succeeding a shock stimulus are shown in panel (a) and mean response times are shown in panel (b) in the random (black squares) and performance-contingent (unfilled circles) shock training groups. The vertical grey line in each figure represents the point at which a shock stimulus was presented. Error bars are between-subjects standard errors.

For our accuracy measure no main effects were observed between our shock conditions and trial number, nor were there any interactions between these factors (all *p*-values > .05). For response times, no main effects of either shock condition or trial number were observed (*p*-values > .05). Although we do find a significant interaction between shock condition and trial number, F(4, 99) = 3.313, *p* = .016, we find no evidence for any time-point specific pair-wise differences in response times between shock condition following Bonferonni adjustment (all *p*-values > .05).

### State Anxiety

We employed the self-report State Trait Anxiety Inventory (STAI) (Spielberger, 1983) measure, using the ‘state’ component response scores as a proxy for participant state anxiety/stress in each of our three shock conditions. These were measured immediately prior to, and following each testing session (pre- and post-test respectively) (see Methods for details). We applied linear mixed-effects modelling to evaluate the effects of shock condition, training session as well as the effect of performing the task on each given day of testing (pre- vs post-testing), with shock condition, training session and daily pre/post-testing as fixed factors and participant as a random factor.

**Figure 9.**
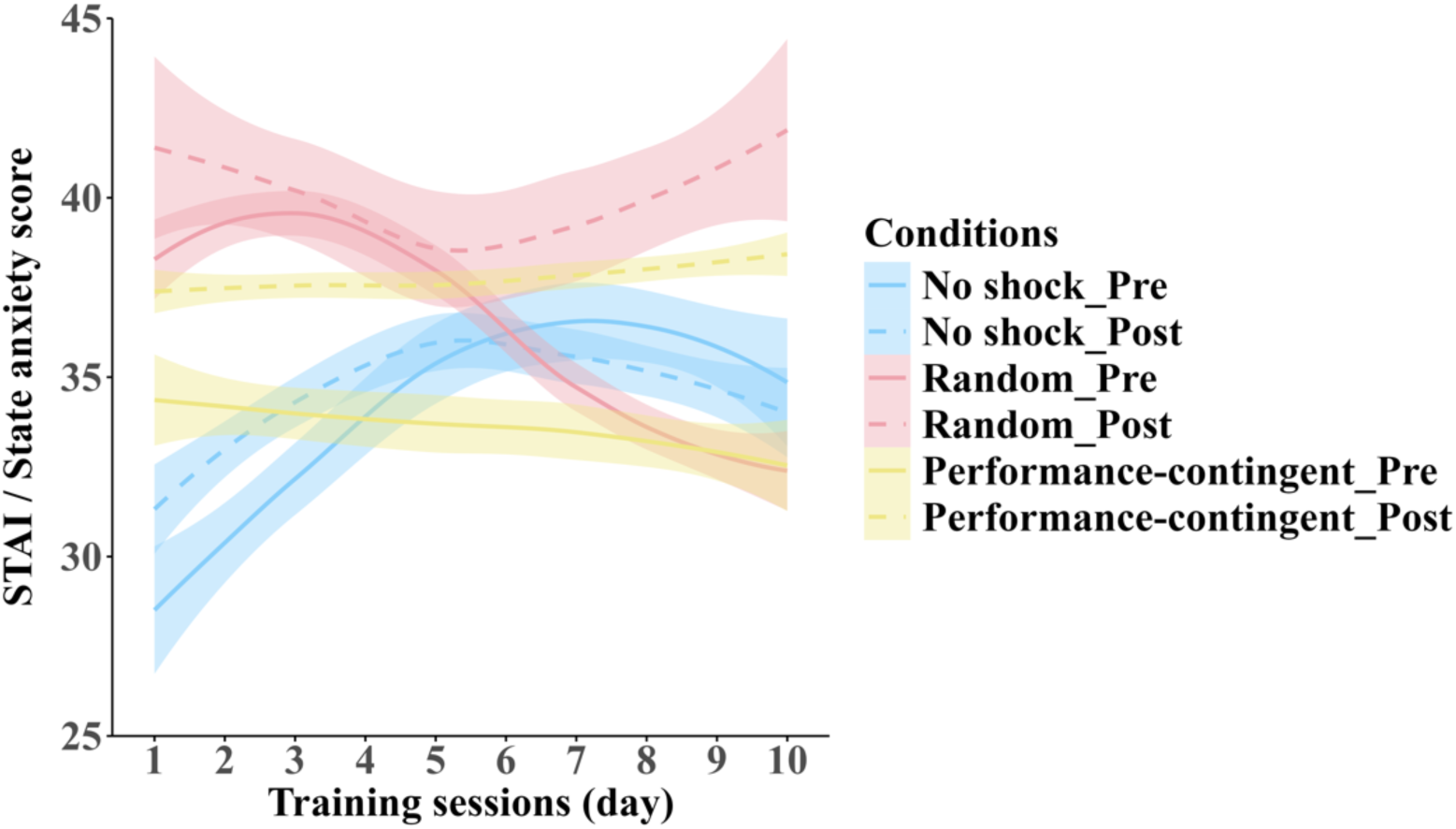
Anxiety before and after training across groups. *Note:* Relationship between average STAI state anxiety subscale scores obtained across training sessions for each shock-condition training group: no shock (blue), random shock (red) and performance-contingent (yellow); measured immediately prior to (solid lines) and following (dashed lines) each training session. Shaded regions represent between-subject standard errors.

We find no evidence for a main effect of shock condition, *F*(2, 12.99) = 0.186, *p* = .832, or training session, *F*(9, 245.01) = 0.326, *p* = .966 on state anxiety. However, a significant main effect of time relative to training session (pre vs post) was observed, *F*(1, 345.00) = 22.032, *p* = .0001, with higher levels of self-reported anxiety observed after daily training than before. Additionally, a significant interaction between the effects of shock condition and time relative to daily training was observed, *F*(2, 245.00) = 3.359, *p* = .036. No other interactions were observed, Shock condition x Training session, *F*(18, 245.00) = 1.167, *p* = .288; Training session x daily training, *F*(9, 245.00) = 0.341, *p* = .960; Shock condition x training session x daily training, *F*(18, 245.00) = 0.601, *p* = .897. To identify the driver(s) of this interaction between shock condition and daily training we ran three within-subjects pairwise comparisons of pre- vs post-test state anxiety scores within each shock condition. While there was no pre- vs. post-testing differences in the *no-shock* condition, *t*(245) = -0.500, *p* = .996, state anxiety scores were on average significantly greater after testing than before testing for *performance-contingent shock* condition, *t*(245) = -4.386, *p* < .001, and the *random shock* condition, *t*(245) = -3.073, *p* = .028.

## Discussion

This study addresses two principal inquiries related to visual processing and perceptual learning. Firstly, we explore whether the human visual system’s capacity to integrate information across the visual field is enhanced through training. This investigation involves manipulating the number of Gabor elements to be averaged (set-size) and subjecting participants to ten consecutive days of training in an orientation averaging task. Secondly, we investigated the influence of acute stress and positive punishment on the rate of perceptual learning in this task.

Several new findings are reported in relation to the first inquiry. Orientation averaging performance, measured in terms of accuracy and response times, consistently degrades with increasing set-size. It is worth noting that we find no differences in the effect of our various shock conditions on the rates at which accuracy and response times degrade with set-size. This reduction in performance with increased set size implies a fundamental constraint on the visual system’s ability to efficiently process and integrate greater amounts of local orientation information.

Notably, training induces significant monotonic improvements in performance across successive testing days, aligning with prior research [19]. Importantly, no evidence of an interaction between set-size and training effects was observed. This indicates that the amount of information the visual system is able to functionally integrate across the visual field to accomplish orientation averaging is not affected by training. Assuming that the observed training-related enhancements in behavioural performance stem from improved sampling efficiency [19], our observation that the *slope* of performance degradation with set-size did not change with training implies that the effects of training did not operate at an ‘early’ encoding stage prior to the integration of local orientation signals. Instead, our results suggest that training exerted improvements in sampling efficiency by increasing signal-to-noise at a ‘late’ post-integration, possibly decisional, stage of processing [20].

Our set-size manipulation includes several distinguishing factors compared to prior orientation averaging studies. A primary distinction is the statistical distribution of Gabor orientations. In classical orientation averaging research, local stimulus features are typically sampled randomly from a global population distribution that follows a Gaussian (normal) distribution with a specified mean and bandwidth. This random sampling can lead to trial-by-trial variations in the mean orientation, meaning that the ‘correct’ response (left or right tilt) may not always align with the physical global stimulus orientation. This probabilistic variation in averaging accuracy has led previous studies to find that performance in global orientation averaging is either unaffected or improves as set-size increases [46, 47].

To avoid these trial-by-trial variations in mean orientation, we adopted a different approach by adjusting the orientation of a single Gabor element per trial to maintain a consistent average global Gabor orientation distribution across trials. However, this method may introduce a distribution that departs from Gaussian characteristics on any given trial, potentially encouraging participants to rely on a "max rule" strategy—where they focus on local deviants rather than averaging orientations across the set. This strategy may contribute to the significant monotonic declines in performance observed with increasing set-size. Future studies could consider using an orientation sampling algorithm that maintains Gaussian-distributed samples with a consistent mean orientation, ensuring more uniform trial structures.

An additional factor that could explain the observed monotonic performance decline with increasing set-size is the phenomenon of visual crowding. Visual crowding refers to impaired target identification when objects are surrounded by featurally similar nearby clutter, reducing perceptual clarity [48-50]. In our paradigm, increasing the set-size of Gabor arrays inherently increased visual density, thus decreasing the spatial separation between adjacent elements. This spatial dependency mimics crowding effects and could lead to declines in performance as set-size grows, as local orientation information becomes harder to isolate amid denser arrays [51]. There are two reasons to believe that the observed performance degradation with increasing set-size is not attributable to visual crowding. First, the distance between adjacent Gabors in our experiment exceeds the spatial extent within which crowding typically occurs, as defined by Bouma’s Law. According to Bouma’s Law, the maximum extent of crowding-related interference zones is roughly half of the target’s visual eccentricity [48, 52]. For observers fixating at the centre of the Gabor array, this limit would be approximately 12.5 degrees, which is larger than the smallest spatial separation in our experiment (set-size of 8, approximate separation 20 d.v.a.). Moreover, because participants were permitted to move their eyes throughout each trial, it is likely that crowding effects were further minimized, as crowding dependencies typically increase with greater eccentricity. Given these factors, it is unlikely that our set-size effects are attributable to visual crowding.

It should be noted that participants in our orientation averaging paradigm were permitted to move their eyes freely, rather than fixating on stimuli at a set eccentricity. Consequently, generalizing our findings to previous work with fixed eccentricities should be done with caution, as eye movement flexibility may influence orientation averaging performance.

It is worth noting that perceptual learning has been found to improve orientation discrimination performance in an orientation-specific manner [53] and to transfer across visual field locations [54]. Whilst it is conceivable that our training-related improvements in orientation-averaging performance may reflect improved local signal-to-noise in early orientation-selective neural channels this seems unlikely as the Gabor patches employed in our study varied randomly from trial to trial not only in their visual locations (polar angles) but also their local orientations. That said, [55] have shown (using Vernier and Landolt C acuity tasks) that perceptual learning transfers to untrained locations and orientations when training includes exogenous attentional cues. This suggests a causal role for attention in the generalisation of perceptual learning across the visual field and feature space. Considering the pivotal role attention plays in the generalization of perceptual learning, coupled with the lack of interaction between set-size and training effects, we posit that our training regimen likely impacted attentional efficiency at a post-integration and/or decisional stage of processing.

Our second major inquiry concerns the effect(s) of acute stress, as operationalised by the presence of an electrodermal shock, on perceptual learning. Whilst we didn’t observe any significant differences in performance between shock conditions on any particular day of testing, we did observe an interaction between training session and shock condition for response times suggesting that the shock conditions influence the improves the rate at which response times improve with training. Of particular significance are the faster response times in the shock conditions as training continued and with greater set sizes.

Noteworthy differences between our shock conditions emerged in terms of *when* subsequent improvements occurred during the training period. Robust improvements in response times were notably delayed until days 4 and 5 in the no-shock and random-shock conditions respectively, contrasting with the performance-contingent shock condition where robust improvements were evident from day 3. Curiously, robust improvement in response times continued to be particularly pronounced up to day 7 in the random shock condition. This temporal variation suggests that our various shock conditions may have exerted differential influences on the urgency of responding during perceptual learning.

Stress, often linked to aversive environments, introduces a complex factor into perceptual learning. Elevated blood cortisol, a physiological response to stress, is associated with changes in cognitive and attentional functioning which inhibits perceptual learning in some tasks [29, 32]. While no study has previously explored the effects of positive punishment or stress on orientation-averaging performance, other studies have shown evidence for benefits of punishment on other visual perception judgements [34]. In the present study, stress responses were assessed using the self-report STAI State Anxiety subscale [38]. Our findings indicate a consistent elevation in state anxiety levels following training, amongst participants exposed to the performance-contingent shock condition. Interpreting this outcome within the context of shock condition group effects on the rate of orientation-averaging perceptual learning, it is suggested that the observed reduction in response times associated with the performance-contingent shock condition may, in part, be linked to the heightened subjective stress experienced in this condition. In contrast, our findings from subjective state anxiety assessments do not provide clear insights into the restricted perceptual learning observed in the accuracy data of participants exposed to random-shock. Crucially, our analysis, which includes all participant responses to maximize statistical power, fails to reveal significant correlations between state anxiety scores and behavioural performance. This implies that our STAI measure inadequately predicts orientation averaging performance, possibly because the STAI may lack adequate sensitivity. Future inquiries employing a larger participant sample and potentially more refined and objective biophysical stress measures, such as saliva cortisol [56] or pupillometry [57], are warranted to elucidate our understanding of the potential effects of stress on orientation averaging performance.

While the current study did not find evidence for an effect of shock on orientation averaging, this is not because orientation averaging is impervious to the effects of external factors. An unexpected aspect of our results is the effect of the health-bar, visible throughout all training sessions and in all shock conditions. We find that orientation averaging accuracy varied according to the health-bar status in all shock conditions, with accuracy improving with increasing levels of visually depicted ‘health’. This is surprising, given that the health-bar was only consequential in the performance-contingent shock condition (predicting the presentation of the shock when the health bar reached zero). In the no-shock and random-shock conditions, the visual representation of health only served to inform participants about accuracy, decreasing with each incorrect response but also resetting to full health following an incorrect response when health reached zero, across trials with no additional consequences. Nevertheless, this visual feedback significantly influenced participant accuracy, with no effect on response times. Interestingly, our time-course analysis revealed that the impact of the five trials immediately preceding and following a shock, under both performance-contingent and random-shock conditions, exhibited no statistically significant differences in performance. This suggests that the observed health-bar effects are not driven by shock anticipation in the performance-contingent condition.

One potential explanation for the unexpected influence of visual performance health-bar feedback on orientation averaging accuracy across different shock conditions is rooted in the literature on attention and perceptual learning. Attention is known to play a crucial role in shaping perceptual learning outcomes, and visual feedback has been shown to modulate attentional mechanisms [58-60]. The health-bar, serving as a continuous visual cue throughout training sessions, may have inadvertently influenced participants’ attentional focus, leading to enhanced orientation averaging accuracy. Feedback can serve as a potent tool for guiding attention and reinforcing learning [61, 62] but see [63]. In this context, the visual representation of ‘health’ could have acted as a motivational cue, encouraging participants to allocate and sustain attention more effectively during the task, thereby improving accuracy in an effort to maintain high health [33]. Finally, the lack of observed effects on response times suggests that the impact of visual feedback on accuracy is unlikely to due to a response-accuracy trade-off.

Recently, virtual reality has been adopted as a vision science tool [64, 65]. Our investigation stands as one of the pioneering studies to utilize virtual reality in the examination of fundamental psychophysics. It is essential to clarify that the first-person shooter task utilized in this study functions solely as a tool for extracting behavioural performance and is not inherently tied to our psychophysical task. In other words, its role is to operationalize psychophysical responses. While a comprehensive validation of our first-person virtual reality paradigm as a tool for investigating fundamental psychophysical performance necessitates a direct comparison with performance using conventional screen presentation (usually with a uniformly coloured background) in conjunction with a standard 2AFC button-pressing task, the pronounced systematic relationship observed in our results attests to the validity of our approach as a psychophysical tool. In consideration of our findings, the virtual reality first-person shooter task holds promise for broader applications in cognitive and vision science research. The task’s immersive nature could be leveraged to investigate various aspects of perceptual and cognitive processes, such as attention, decision-making, and spatial cognition. Furthermore, its potential to simulate real-world scenarios may offer valuable insights into how individuals respond to visual stimuli in ecologically valid settings.

## Conclusion

Our findings contribute valuable insights into the mechanisms of perceptual learning and the impact of aversive stimuli on visual processing. The training-induced improvements in orientation averaging performance, particularly the lack of interaction with set-size, suggest that the enhancements operate at a post-integration and/or decisional stage of processing. The distinctive effects of aversive shock conditions on perceptual learning underscore the complex interplay between stress, attention, and motivational processes on performance. The unexpected influence of visual feedback on accuracy highlights that the visual averaging task is not impervious to the effects on contextual information and warrants further investigation into the potential importance of attention and possibly motivation, in perceptual learning.

Additionally, our use of virtual reality, particularly the first-person shooter task, represents an innovative approach in the study of fundamental psychophysics. The task’s potential for broader applications in cognitive and vision science research opens avenues for investigating attention, decision-making, and spatial cognition in ecologically valid settings. While our study sheds light on the computational principles underlying perceptual learning in our orientation averaging task, further research with larger participant samples and refined stress measures is recommended to deepen our understanding of the effects of stress on learning and perceptual performance.

## Author contributions statement

JC, GW & WHF wrote the initial manuscript

JC & GW designed the experiment and experimental protocol

JC supervised the experimental data collection

JC, GW, WHF & LC wrote the initial manuscript

YL ran the experimental analyses and conducted the data visualisation

All authors reviewed the manuscript.

